# Dopamine in the songbird auditory cortex shapes auditory preference

**DOI:** 10.1101/761783

**Authors:** Helena J. Barr, Erin M. Wall, Sarah C. Woolley

## Abstract

In vocal communication, vocal signals can provide listeners with information and also elicit motivated responses. Auditory cortical and mesolimbic reward circuits are often considered to have distinct roles in these processes, with auditory cortical circuits responsible for detecting and discriminating sounds and mesolimbic circuits ascribing salience and modulating preference for those sounds. Here, we investigated whether dopamine within auditory cortical circuits themselves can shape the incentive salience of a vocal signal. Using female zebra finches, who show natural preferences for vocal signals produced by males (‘songs’), we found that pairing passive song playback with pharmacological manipulations of dopamine in the secondary auditory cortex drives changes to song preferences. Plasticity of song preferences by dopamine lasted for at least one week and was not influenced by norepinephrine manipulations. These data suggest that dopamine acting directly in sensory processing areas can shape the incentive salience of communication signals.

## INTRODUCTION

Vocal signals are critical for survival and reproduction in a range of species. Receivers can extract substantial information from vocal signals about the identity, species, or motivation of the signaller and make mate-choice and other social decisions based on the incentive salience of the signal. However, there is growing consensus that receivers, and their auditory systems, are not passive filters, but rather they dynamically encode acoustic stimuli.^1,2^ Consequently, a signal’s salience may not be an inherent component of the signal, but instead be determined by the individual receiver’s internal state and experience^3,4^. For example, in fish, frogs, and birds, reproductive status, acting through changes in steroid hormones and neuromodulators, can influence auditory responses and the processing of mating calls^5–7^. Similarly, maternal experience and reproductive status dramatically shape the way that female rodents respond to pup calls, in part due to neuromodulatory shaping of auditory responses^8–10^. Thus, the response to vocal communication signals depends not only on the signal itself, but also on the ascribed salience of those signals to an individual receiver.

Dopamine (DA) is a key modulator for ascribing incentive salience to stimuli, providing the brain with information on which sensory stimuli are relevant or important^11–13^. Dopamine neurons in the ventral tegmental area (VTA) respond to reward-related stimuli in learning tasks across sensory domains^14,15^. Moreover, dopaminergic projections from the VTA to regions like the nucleus accumbens have been found to influence a wide range of motivated behaviors^13,14,16^. For example, DA acting within the nucleus accumbens can shape behavioral responses and preferences for particular stimuli, including social stimuli. In male prairie voles, injections of DA agonists into the nucleus accumbens induces partner preference^17,18^. Thus, dopaminergic activity can serve to modulate behavioral responses to communication signals.

However, little is known about how DA can act on areas outside the traditional mesolimbic pathway to influence incentive salience for sensory stimuli. Recent studies have documented that DA signals from the VTA can shape activity and tuning in the auditory cortex^19,20^ while DA signals in the nucleus accumbens relate to reward value or incentive salience^12^. This has led to the model that DA from the VTA simultaneously acts at the level of the nucleus accumbens to shape preferences and at the level of sensory cortex to correspondingly shape sensory tuning to those stimuli^11,21^. However, it is not known whether DA acting in the sensory cortex itself could drive changes in the salience and incentive value of sensory stimuli.

Here, we investigated the degree to which preferences for particular vocal communication signals can be altered by manipulating neuromodulatory input to the auditory cortex. We studied this in the zebra finch, a species of songbird in which adult females identify individuals and select mates based on their complex, learned vocalizations (‘songs’). We found that pharmacological manipulation of dopaminergic activity in the auditory cortex significantly shaped preferences for song and could reverse preferences for some songs over others. These data suggest that DA can act directly in sensory processing areas to shape the incentive salience of and preferences for stimuli.

## RESULTS

### Dopamine neurons in the caudal VTA show greater responses to preferred songs

We first quantified female preferences for songs using a two-choice operant assay (Fig. 1A)^22,23^. In this assay, female zebra finches were provided two strings, each of which activated the playback of a song from a single male zebra finch when pulled (e.g., Male A for one string, Male B for the other string). For each of the five song pairs tested, females showed significant song preferences for one of the songs of the pair (p<0.01 for all; Fig. 1B; see Methods). For four of the five pairs, there was variation across females in which song was preferred (e.g. some females preferred Male A, others Male B; Fig. 1C). However, for one of the pairs of songs (Male J and Male K), females consistently preferred the song of one male of the pair over the song of the other male (Fig. 1B; t_(16)_=4.29, p=0.0006).

**Figure 1.**
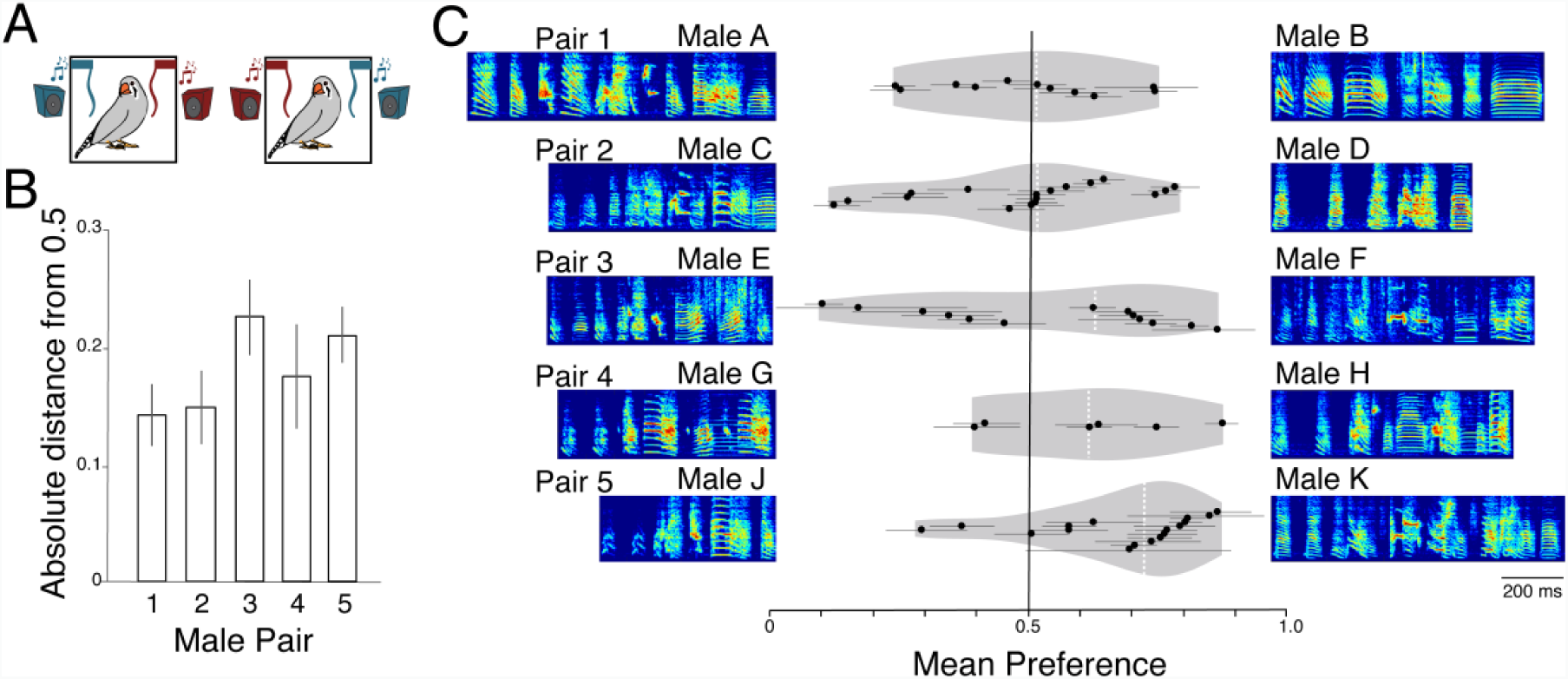
Females song preferences revealed by string pull assay. A) Females were tested in a string pull assay where each string triggered playback of a song of a particular male. Contingencies were reversed halfway through testing to control for side bias. B) The absolute distance in preference score from 0.5 (which corresponds to a lack of bias for one specific song) was significantly greater than zero for all five pairs, indicating that females showed preferences for the song of one male over another. C) While each individual female showed significant preferences for one song within all pairs of male songs, there was significant individual variation in which male was preferred for all but one of the pairs. Violin plots (gray shading indicating the probability density) show the responses of females to pairs of songs, indicated by spectrograms of the song motifs at each end of the plots. Points are the responses of individual females with horizontal lines from the point indicating bootstrapped confidence intervals. Vertical white dashed lines indicate the median of preferences across all females. Motifs are labeled with an ID for each male and ordered by male pair as in B. Only for the bottom pair of males (Pair 5: Male J and Male K) did female preference differ significantly from 0.5, indicating consistent preferences for Male K across females.

We leveraged the fact that a majority of females clearly preferred one of the two males in Pair 5 (Fig. 1C) to investigate whether catecholaminergic neurons in the midbrain and hindbrain differentially respond to songs with different degrees of incentive salience, i.e. preferred versus less preferred songs. We calculated the percent of tyrosine hydroxylase (TH) neurons that expressed the immediate early gene FOS in birds that heard either the preferred or less-preferred song, or were left in silence. We found significant variation in responses across the five brain regions we measured (F_(8,109)_=14.212, p<0.0001; Fig. 2). The caudal ventral tegmental area (cVTA) was the only region to differentially respond to preferred vs. less-preferred song. In particular, a greater percentage of DA neurons in the cVTA expressed FOS following playback of preferred song than following playback of the less-preferred song (p=0.0013) or silence (p=0.0002). In contrast, while more TH neurons in the locus coeruleus (LC) and periaqueductal gray (PAG) expressed FOS in response to hearing songs (p<0.0001 for each), these neurons responded similarly to preferred and less-preferred songs (PAG: p=0.4943; LC: p=0.0630). These data support previous work that finds that catecholamine-synthesizing neurons in the midbrain and hindbrain respond to playbacks of social signals^24^. In addition, these data highlight that dopaminergic neurons in the VTA, but not in other catecholamine-producing cells in the midbrain and hindbrain, are differentially activated by songs with different degrees of incentive salience. Together, this suggests that songs with different incentive values lead to differing amounts of dopamine release in areas downstream to the VTA.

**Figure 2.**
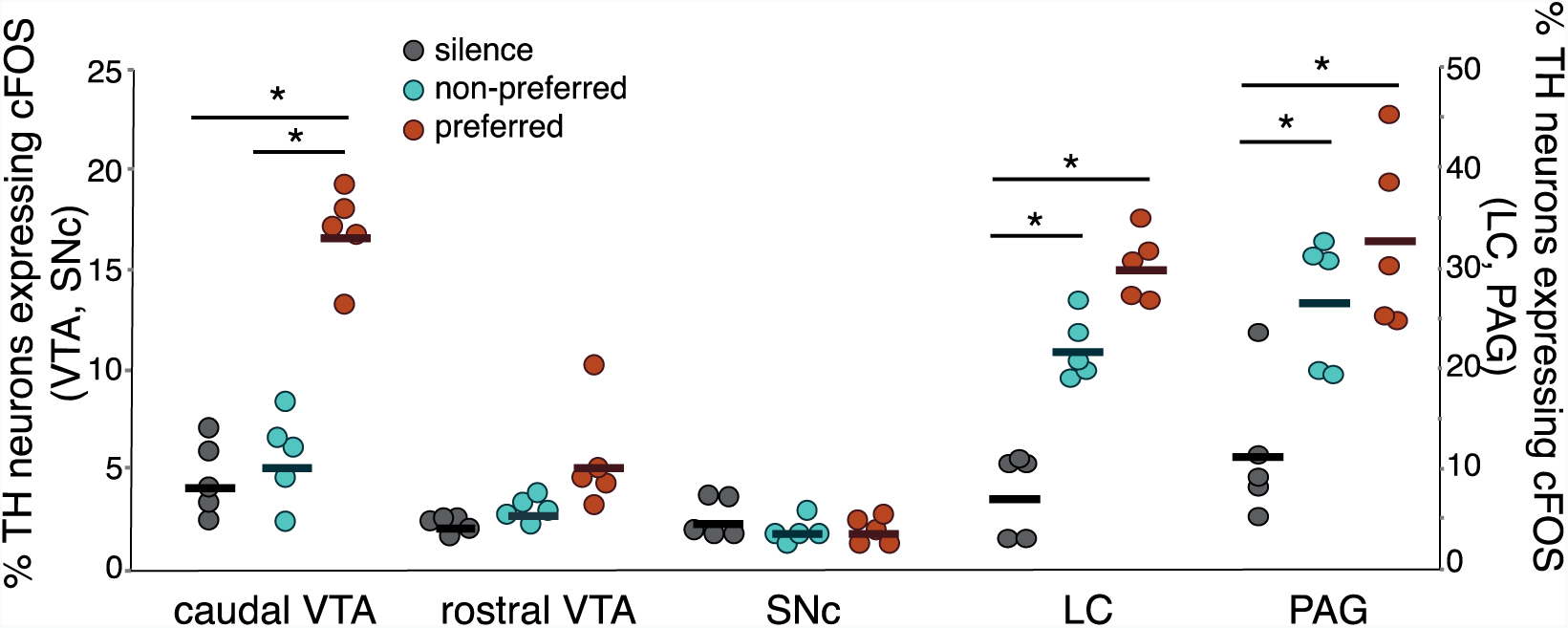
Hearing preferred songs drives FOS expression in dopaminergic neurons of the caudal VTA. The percent of tyrosine hydroxylase (TH) neurons expressing FOS in the ventral tegmental area (VTA) and substantia nigra pars compacta (SNc; left axis) and the locus coeruleus (LC) and periaqueductal gray (PAG; right axis) following playback of preferred songs (orange) or non-preferred songs (teal) and in silent controls. In the caudal VTA, preferred song elicited significantly more FOS expression in TH neurons than either non-preferred song or silence. In the LC and PAG, both preferred and non-preferred songs elicited more FOS in TH neurons, however, there was no significant difference in the percent of TH neurons expressing FOS between preferred and non-preferred songs.

### Pairing song playback with a general dopamine agonist shifts song preference

To investigate whether catecholamine release into the auditory cortex could shape song preferences, we paired infusions of catecholaminergic drugs into the secondary auditory pallium (Fig. 3A and 3B) with passive playback of a male’s song. We targeted the caudomedial nidopallium (NCM), a secondary auditory region important for auditory processing and implicated in auditory memory^25–28^. In particular, after measuring the song preferences of individual females (Day 1), we paired infusions of catecholamine agonists with playback of the less-preferred song and paired infusions of vehicle (5% DMSO in phosphate buffered saline (PBS); see Methods) with playback of the preferred song (Days 2 and 3). Females were then re-tested for song preferences (Day 4; i.e., preference for one song compared to the other song of the pair; same as Day 1; see Methods; Fig. 3B).

**Figure 3.**
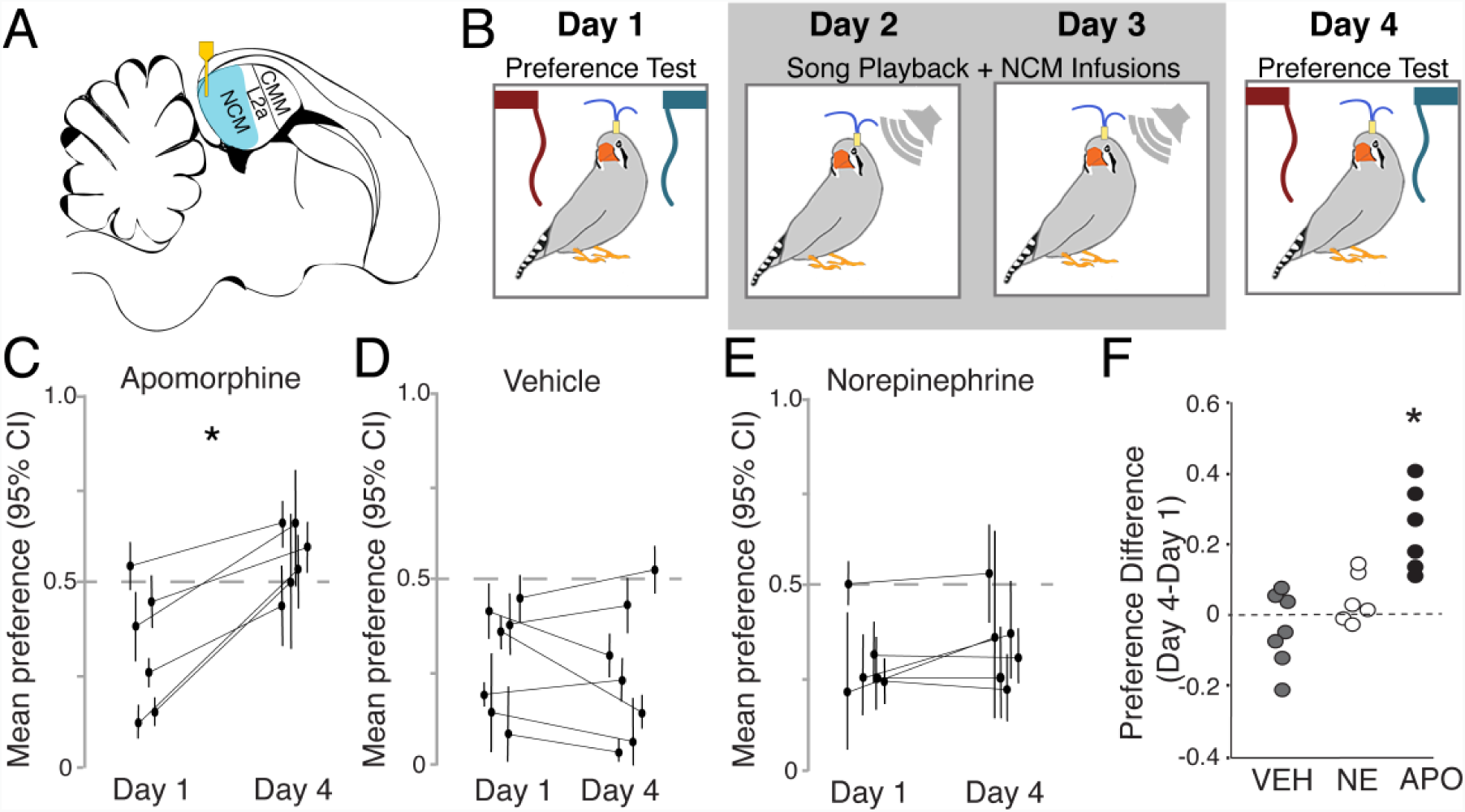
Dopamine in NCM modulates song preference. A) Microdialysis cannulae were implanted into the caudomedial nidopallium (NCM; blue). Parasagittal illustration of the location of the cannulae (yellow) relative to the NCM, the input “layer” of the primary auditory pallium Field L (L2a), and the secondary auditory pallial region the caudomedial mesopallium (CMM). Dorsal is up, caudal is left. B) Diagram of the experimental protocol. Females were tested for preference using a string pull assay on Day 1. On Days 2 and 3 females were infused with either vehicle (VEH) or drug 10 minutes prior to 2-hours of song playback. Drug and song combinations during Days 2 and 3 depended on the experiment (see Methods). In general, for agonist tests, agonists were paired with playback of the less-preferred song and VEH was paired with the preferred song. For control tests, VEH infusions were paired with playback of the less-preferred as well as the preferred songs. On Day 4, females were tested again in the string pull assay. The order of drug and control infusions (Day 2 and Day 3) were randomized across birds and tests. C) Females given the general agonist apomorphine paired with playback of the less-preferred song significantly shifted their preferences toward the originally less-preferred song. Points are the preference for the less-preferred song for individual birds, bars indicate bootstrapped confidence intervals (see Methods). No significant changes in preference from Day 1 to Day 4 were observed in vehicle treated control females(D) or in females given norepinephrine (E). F) The change in preference between Day 1 and Day 4 was significantly higher in females that received apomorphine (APO) than females infused with VEH or norepinephrine (NE).* indicates p<0.05.

We first tested the effects of broad, general catecholaminergic agonists to determine whether DA or norepinephrine (NE) could alter female song preferences (change in preference scores from Day 1 to Day 4). We found that pairing the less-preferred song with the general DA agonist apomorphine (APO) significantly affected female preferences for song (t_(5)_=-5.09, p=0.0038). Specifically, pairing playbacks of the less-preferred song with APO infusions into the NCM led to a significant increase in preference for that song between Day 1 and Day 4 (Fig. 3C). In contrast, song preferences were stable and unchanged between Day 1 and Day 4 in the control condition, when playback of the less-preferred song (as well as to the preferred song) was paired with vehicle (VEH; t_(6)_=1.06, p=0.3306, Fig 3D). Similarly, pairing the less-preferred song with NE in the NCM also did not significantly alter preferences (t_(5)_=-1.52, p=0.1895; Fig. 3E).

To compare directly between drugs, we calculated the change in preference from Day 1 to Day 4 and compared the degree of change between the three treatments (Fig. 3F; see Methods). Overall, there was significant variation across treatments (F_(2, 12.4)_=10.43, p=0.0022), with a greater change in preference for APO than for either NE (p=0.0334) or VEH (p=0.0017). Thus, when coupled with passive song playback, DA, but not NE, in a secondary auditory cortical region modulates song preference. These data indicate that DA in the secondary auditory cortex influences the incentive salience of songs. Given this finding, it is important to reveal the involvement of specific DA receptor subtypes, persistence of the effect, and nature of drug pairing on female auditory preferences.

### D1 receptors participate in shifting song preference

D1-type receptors are highly expressed in the auditory forebrain, including NCM^29^. To investigate the degree to which D1-type receptors are involved in song preferences, we paired song playback with D1 receptor-specific agonists, antagonists, or VEH (see Methods). Like APO, pairing infusions of the D1 receptor-specific agonist SKF81297 into the NCM with playback of the less-preferred song led to a significant increase in the preference for the less-preferred song (t_(5)_=-5.10; p=0.0038; Fig. 4A) such that females no longer demonstrated a significant preference for the previously preferred song. Conversely, pairing playback of the preferred song with infusions of the D1-receptor antagonist (SCH23390) tended to decrease the preference for the preferred song (t_(6)_=-1.84; p=0.1147; Fig. 4B). Both D1-receptor drugs produced greater shifts in preference relative to the vehicle control condition (F_(2,6.62)_=45.31; p=0.0001; Fig. 4C). In particular, the shift in preference was significantly greater for the D1-receptor agonist than for VEH (p=0.0001), and tended to be greater for the D1-antagonist than for VEH (p=0.0541). Together these data indicate that manipulation of D1 receptors in the auditory cortex during song playback can produce substantial changes in preference and highlight the importance of D1 receptors in the auditory forebrain in ascribing incentive salience to stimuli.

**Figure 4.**
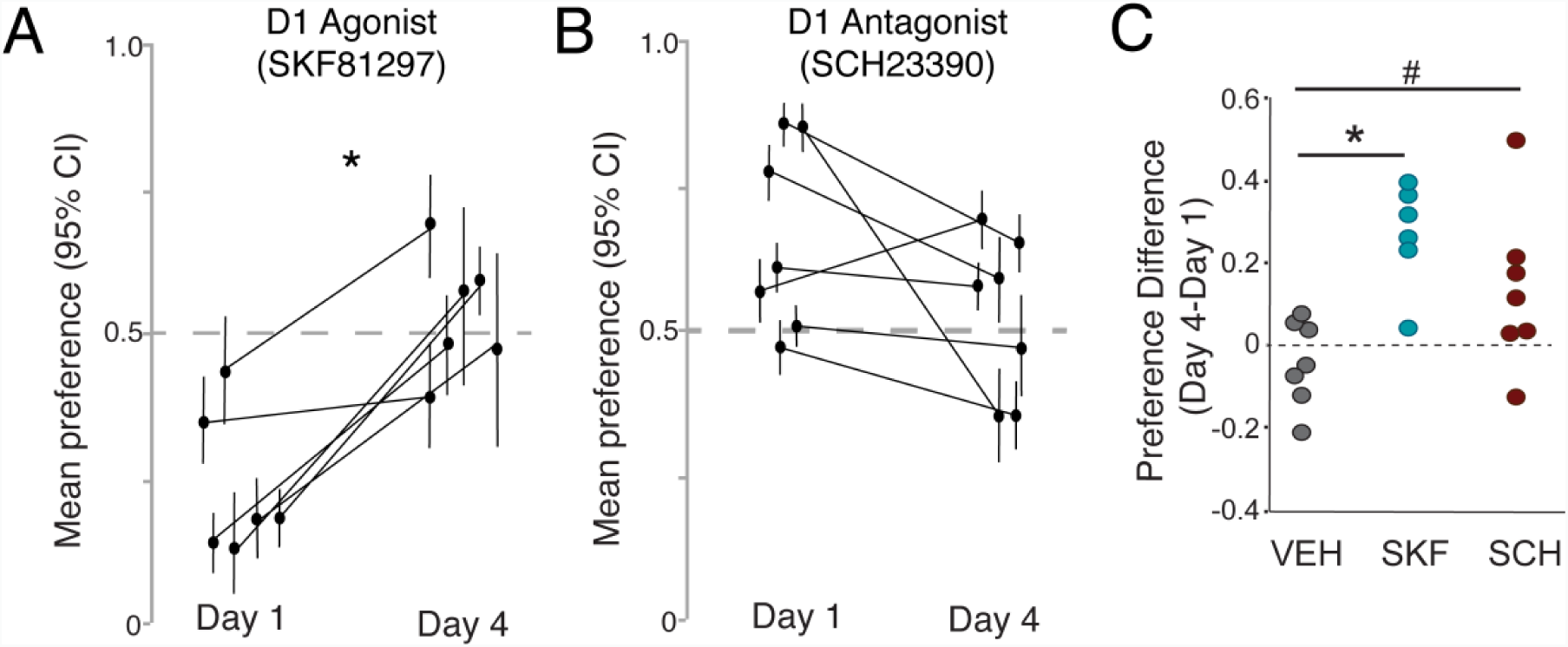
Dopamine D1 receptors in the NCM affect song preference A) Females given the D1 receptor agonist SKF81297 paired with playback of the less-preferred song significantly increased their preferences toward the originally less-preferred song. Points are individual birds, bars indicate bootstrapped confidence intervals (see Methods). No preference is at 0.5 indicated by the dashed line. B) Females given the D1 receptor antagonist SCH23390 paired with playback of the preferred song show a trend toward diminished preference for the preferred song. C) The change in preference between Day 1 and Day 4 was greater in females that received the D1 receptor agonist or antagonist than females infused with VEH.* indicates p<0.05; # indicates p=0.054.

### Song preference shifts are not a consequence of changes in motor behavior

Given the known roles of dopamine in modulating motor behavior and motivation, we investigated whether the drug manipulations had general effects on string pulling behavior overall and the degree to which these changes could have contributed to the shifts in preference. While across all treatments, the total amount of string pulling for either string on Day 4 was lower than that on Day 1 (F_(1, 43.2)_=8.78, p=0.0049), the shifts in preference were not due to the overall changes in string pulling. Although different drugs had different effects on female preferences, the degree to which females changed the amount of string pulling did not vary across drugs (Drug X Day interaction: F_(4,43.2)_=0.49, p=0.7455; Suppl Fig. 1A). Further, the percent change in string pulling did not significantly correlate with changes in preference for any of the conditions (Suppl Fig. 1B; p>0.05 for all conditions). Taken together, these data indicate that the changes in preference were not due to general effects of drug manipulations on motor behavior or the motivation to hear song.

### DA infusions into NCM lead to lasting changes in song preferences

To determine the degree to which manipulation of dopamine in the NCM can lead to lasting changes in preference, we also tested a subset of birds one week following pairing of song playback and the D1 agonist (see Methods). We found that these females continued to showed preferences that were significantly shifted from baseline (Fig. 5A). In particular, females tested one week after pairing of the D1 receptor agonist and the less-preferred song had significantly increased preferences for the previously less-preferred song (t_(5)_=-4.76, p=0.0050). Moreover, the magnitude of the shift in preference was not significantly different from females tested immediately following pairing of the D1 receptor agonist and the less-preferred song (F_(1,10)_=0.05; p=0.8371; Fig. 5B). Thus, pairing of song playback with D1 receptor stimulation can result in lasting changes in song preference.

**Figure 5.**
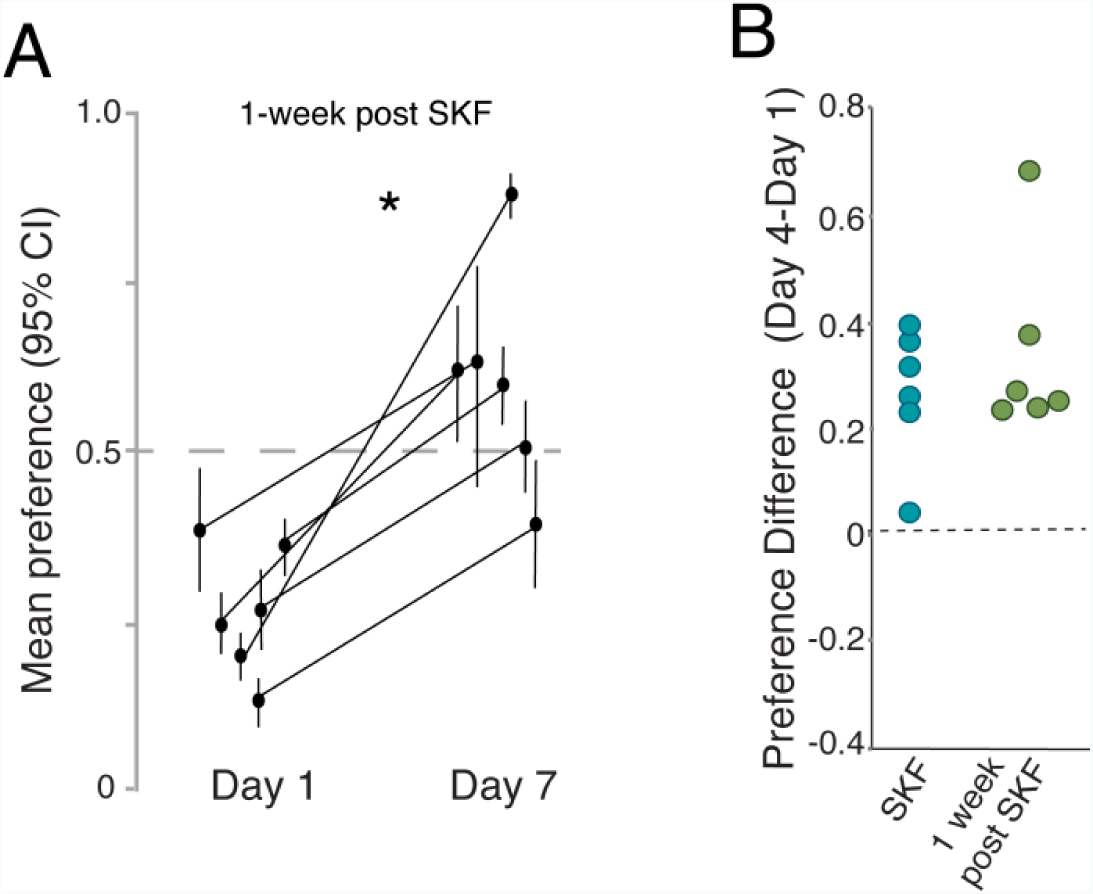
Lasting preference shifts following DA infusions paired with playback. A) Females tested for preference one week following pairing of the D1 receptor agonist with playback of the less-preferred song showed significant increases in preference for the less-preferred song. Points are individual birds, bars indicate bootstrapped confidence intervals (see Methods). No preference is at 0.5 indicated by the dashed line. B) Preference changes induced by D1 agonist paired with playback of the less-preferred song for females tested on Day 4 (“SKF”; teal) did not differ significantly from the preference changes of females tested 1 week later (green). “*” indicates p<0.05.

### Shifts in preference depend on the timing of drug infusion relative to song playback

The effects of VTA stimulation on tonotopic maps in the primary auditory cortex depend on the timing of stimulation relative to sound playback^19^. This suggests the possibility that D1-receptor-mediated changes in preference may require a temporal correspondence between drug infusion and song playback. To investigate the importance of the temporal association between drug infusion and song playback for changes in preference, we uncoupled the timing of drug delivery and song exposure and estimated changes to song preferences (see Methods). Specifically, during the song exposure on either Day 2 and 3, females were infused with the D1 agonist for 2-hrs beginning 15-30 minutes after the termination of playback of the less-preferred song. This uncoupling of drug infusion and song playback did not lead to a significant shift in song preferences (t_(5)_=1.11, p=0.3181; Fig. 6A). Moreover, the change in preference when the D1-receptor agonist was uncoupled from song was significantly less than when the D1-receptor agonist was coupled with song playback (p=0.0033; Fig. 6B), and not significantly different than the lack of preference change following VEH (p=0.8283). Together, these data indicate that the reorganization of song preference requires the temporal coupling of D1-receptor activation and song playback.

**Figure 6.**
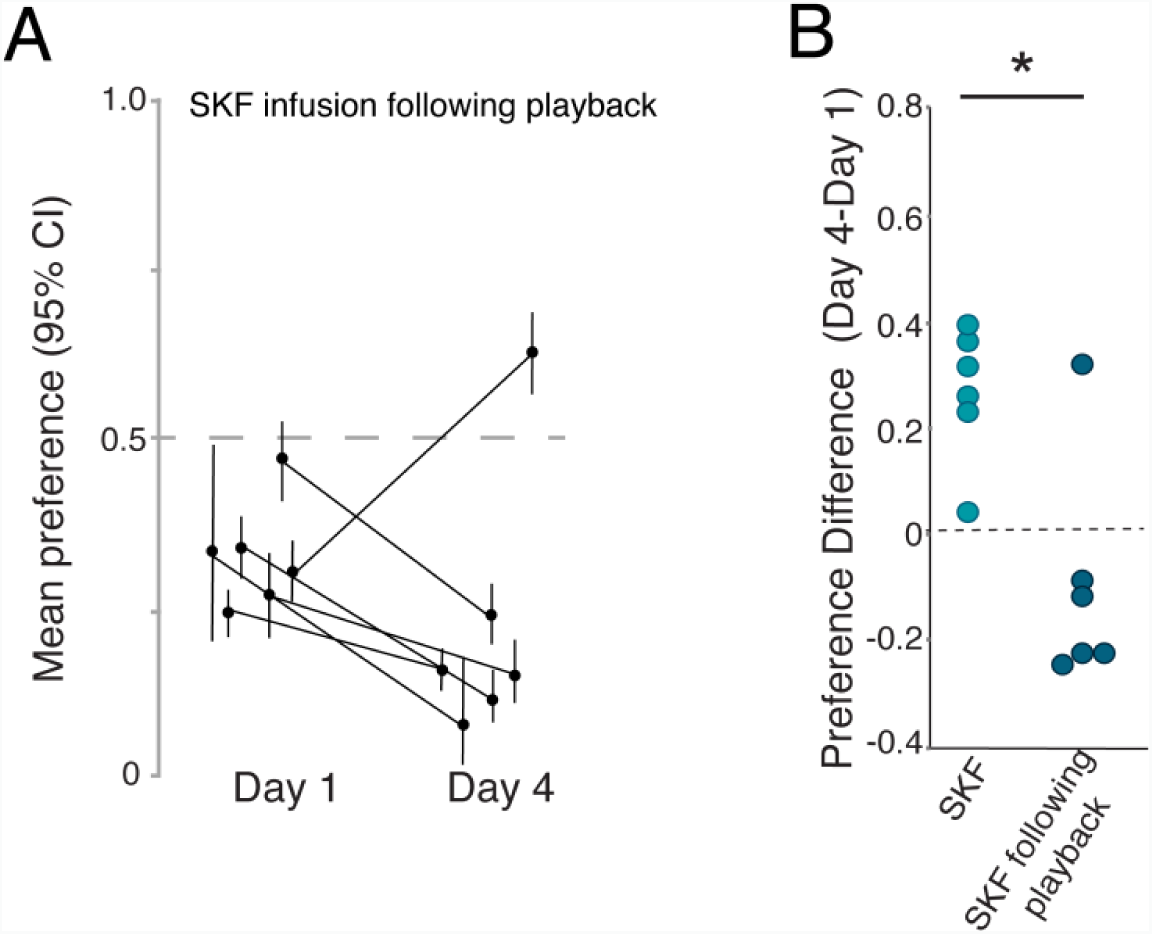
Preference shifts are dependent upon the timing of drug infusion. A) Infusing the D1 receptor agonist into NCM after song playback did not lead to greater preference for the less-preferred song. Points are individual birds, bars indicate bootstrapped confidence intervals (see Methods). B) Comparison of preference changes induced by D1 agonist paired with playback of the less-preferred song for females tested on Day 4 (“SKF”; teal) and of females that received SFK infusions 15-30 minutes following song playback (blue). “*” indicates p<0.05.

## DISCUSSION

In vocal communication systems, receiver preferences for vocal signals can depend not only on the features of the signal but also on the receiver’s individual experience or internal state. The dual tasks of processing a signal’s acoustic features and its incentive value have been postulated to rely on physiological processes within sensory and reinforcement pathways respectively. In particular, auditory cortical circuits are thought to be responsible for detecting and discriminating acoustic signals while the nucleus accumbens and VTA ascribe salience and modulate preference for those signals. In line with this, we found that in female zebra finches preferred songs elicit greater activity in dopamine neurons of the caudal VTA than less-preferred songs. However, we also found that coupling song playback with pharmacological manipulations of dopamine receptors within the auditory cortex itself could alter and, in some cases, fully reverse song preferences. The changes to preference were lasting, as females still displayed the reversed preference one week later. Moreover, the changes to preference were not just a consequence of dopamine-dependent changes in the auditory cortex, since females that heard playback before drug manipulation did not show altered preferences. Taken together, these data indicate that dopaminergic projections to the auditory forebrain may directly modulate behaviorally-relevant auditory preferences and motivate behavior in response to vocal signals.

Pairing DA agonists in the NCM with playback of a less-preferred song resulted in increased preferences for the less-preferred stimulus. Thus, song preferences are not only plastic under the right conditions, but they can be altered by changes in the auditory forebrain. In rats, stimulation of the VTA leads to changes in the inhibitory tone and plasticity in the tonotopic organization of primary auditory cortex (A1)^19,30^. Specifically, VTA stimulation enhances circuit inhibition in A1, thereby increasing the auditory-evoked firing precision of A1 neurons^30^. Moreover, pairing VTA stimulation with playback of a tone leads to expanded representation of that tone in A1^19^. Such targeted changes to A1 firing precision and tonotopic organization are associated with increased ability to discriminate sounds^31–33^. However, while previous studies have shown that VTA stimulation leads to auditory cortical plasticity and improved discrimination, changes just to the representation of a sound have not been hypothesized to result in changes in preferences. Indeed, in our study, birds were not simply discriminating between two stimuli, but were pulling strings to hear one song over another; therefore, their string pulling behavior provided a read-out of their motivation to hear a particular stimulus. Thus, our data indicate that increasing dopaminergic activity in the auditory forebrain could lead not only to an increase in the signal-to-noise ratio, as seen in rodents, but also to a change in the motivation to hear the song or the pleasure derived from it.

At the same time, we found that preference for a preferred song could be diminished following pairing of playback with infusion of a D1-receptor antagonist. Thus, our data support the possibility that dopamine is released into the NCM in response to preferred songs and this release may be important for the sustained preference for that song. These data dovetail with previous work demonstrating that, in female sparrows, hearing conspecific song (versus silence) leads to increases in the expression of phosphorylated tyrosine hydroxylase, a marker of dopamine synthesis, in the NCM and CMM^34^. Together with the lower expression of FOS in response to less-preferred songs, this indicates that least preferred songs may elicit lower levels of dopamine release. Future studies using online methods to measure local changes in dopamine release^35–37^ in the auditory cortex in response to songs of different perceived quality will provide needed data to clarify this relationship.

Dopamine release in the striatum and the cortex leads to plasticity through both long-term potentiation (LTP) and depression (LTD)^38–42^. For example, in the songbird basal ganglia nucleus Area X, induction of LTP requires activation of both NMDA and D1 receptors^38^, while in the rat prefrontal cortex dopamine lowers the threshold for LTD^42^. The effects of dopamine receptor activation on synaptic plasticity in primary sensory areas have not been directly addressed, however, one interesting possibility is that, through similar synaptic mechanisms, pairing of song and dopamine stimulation may lead to changes in the encoding of auditory objects that could lead to enhanced memory of the song^43,44^. The NCM has been implicated in auditory memory for songs, including memories of the song of the tutor as well as a mate^26,45,46^. Moreover, female songbirds show long-lasting memory for the songs of specific familiar individuals, including songs of the tutor and the mate^26,47–49^. In males, NCM neurons become more selective for the tutor song following tutoring^27^. Whether similar processes occur in females, and the degree to which they are dopamine-dependent is unknown. Future investigation of whether dopamine in the NCM not only modulates song preference but also leads to the formation of auditory memories for especially salient or preferred songs, such as the mate’s song, will provide needed and novel insight into the function of the NCM in auditory perception.

Attribution of social salience to acoustic signals is a critical step in auditory processing for vocal communicators. Our data extend existing knowledge about catecholamine function in the auditory cortex^15,19,30,50,51^. Moreover, our results implicate the auditory cortex in the shaping of auditory preferences. Thus, dopamine-dependent changes in the auditory cortex may not only increase representational distinctiveness, heighten signal-to-noise, and improve discrimination ability, as has been seen in studies on rodents, but these changes can lead directly to a change in the incentive salience of the sound.

## METHODS

### Animals

All zebra finch females used in this study (N=36, >90 days post-hatch) were raised with both parents and all siblings until 60 days of age. Thereafter, they were housed in same-sex group cages in a colony, and thus were acoustically exposed to the vocalizations of both males and females. Females were maintained on a 14-hour light, 10-hour dark schedule with *ad libitum* access to seed, water, and grit. Lettuce and egg supplements were provided once per week. Bird care and procedures followed all Canadian Council on Animal Care guidelines and were approved by the Animal Care Committee of McGill University.

### Drugs

All drugs were dissolved in DMSO and diluted in sterile phosphate buffered saline (PBS) such that the final concentration of DMSO was 5%. We used three drugs that target dopamine receptors: Apomorphine (APO, a general dopamine agonist; 3.3mM; Tocris Bioscience, Minneapolis, MN), SKF-81297 (1mM; a selective D1-receptor agonist; Sigma Aldrich, Oakville, ON), SCH-23390 (1mM, a selective D1-receptor antagonist, Tocris Bioscience, Minneapolis, MN). We also assessed the effects of norepinephrine (NE; 1 mM; Sigma Aldrich, Oakville, ON) on song preferences. Drug concentrations were determined based on the literature and personal communication^52–56^. For all tests, PBS containing 5% DMSO was used as the control vehicle solution (VEH).

### Surgery

To manipulate catecholamine levels in NCM, females were bilaterally implanted with microdialysis guide cannulae targeted at the NCM (Fig 3A). At least 30 minutes prior to surgical procedures, females were given an analgesic (Metacam, company) and deprived of food and water. At the start of surgery, females received an intramuscular injection of ketamine (0.04mg/g) and midazolam (0.0015mg/g) for anesthetic induction and then fitted into a stereotaxic apparatus (Leica) with a fixed beak angle of 45 degrees. Once birds were placed into the stereotaxic apparatus, anesthesia was maintained on 0-2% isoflurane vapor for the duration of the surgery. Guide cannulae containing dummy probes (CMA/7, CMA Microdialysis, Stockholm, Sweden) were implanted bilaterally in the NCM (from the caudal Y-sinus: 50 μm rostral, 50 μm lateral, 150 μm deep) through small windows in both layers of skull and secured in place using epoxy and dental cement. Following surgery, all females were housed individually and given at least a week to recover before beginning retrodialysis and behavior testing.

### Reverse microdialysis

Females were fitted with microdialysis probes into the guide cannulae and infused with VEH at least 24 hours before the start of the experiment to allow for habituation. Solutions were retrodialyzed into the NCM using untethered microdialysis probes (CMA Microdialysis, Kista, Sweden; pore size 6,000 Daltons). Specifically, probe input and output tubing were trimmed to 3-4 cm and fitted with connectors and custom-made stoppers. Outside of the experimental period, females were infused every 12 hours with VEH using a syringe pump (Harvard Apparatus, Holliston, MA; 10 μl/min for 4 minutes). On infusion days (Days 2 and 3), tubing was filled using the syringe pump with 40 ul of drug or VEH via the input tubing. Following song playback exposure, tubing was flushed with VEH (10 μl/min for 4 minutes) then filled with 40 ul of VEH (10 μl/min for 4 minutes).

### Preference testing

For the duration of testing, females were individually housed in sound-attenuating chambers (TRA Acoustics, Cornwall, Ontario) inside cages equipped to test song preferences with a string-pull assay. Specifically, cages contained two Cherry 1g levers, each with a piece of a burlap string attached. Levers were connected to a computer via a connector block (National Instruments). Sound Analysis Pro software was used to record string pulls and playback songs^22,57^. During song preference testing, levers were activated so that each string, when pulled, triggered the playback of one male’s song. For example, String 1 triggered the playback of Male A’s song, and String 2 triggered the playback of Male B’s song (Fig 3B). Preference tests consisted of two 2-hour sessions. Females were required to pull each string a minimum of three times to initiate each session. For the second session, the song triggered by each string was switched (i.e., contingencies reversed; for example, now String 1 triggered Male B’s song, and String 2 triggered Male A’s song) to control for place/string preference. Following the switch in contingencies, the second session began once the female had pulled each string a minimum of three times. At the end of the second session, all strings were removed from the cage.

### Experimental Design

The experiment followed a four-day schedule. On Day 1, a female’s preference between two male songs was tested using the string pull assay, allowing us to identify the “preferred song” and “less-preferred song.” On Days 2 and 3, females received retrodialysis of drug or VEH (one treatment per day) during two hours of passive exposure to the songs of one of the males. The order of song exposure (preferred vs. less-preferred songs) on days 2 and 3 was randomized within each female across multiple experiments as well as between females. For tests using DA and NE agonists, we paired playback of the less-preferred song with infusion of either a DA receptor agonist (APO, SKF-81297) or NE, and paired playback of the preferred song with infusion of VEH. All infusions occurred 10 to 30 minutes prior to the beginning of playbacks, and all drugs were washed out (10 μl/min for 4 minutes) within 30 minutes following the end of playbacks. On Day 4, the female’s preference was retested (Fig 3C). For tests assessing whether DA antagonism could decrease preference for the preferred song, we paired playback of the preferred song with infusion of SCH-23390 and playback of the less-preferred song with VEH. In the control test, VEH was infused prior to playback of both the preferred and less-preferred songs. Females each underwent multiple experiments, with different pairs of male songs for each drug manipulation. The pair of males paired with each drug manipulation and the order of drug manipulation was randomized across females.

We also tested whether DA manipulation was necessary during the song playback in order to affect preferences. In a separate set of experiments, the DA agonist SKF81297 was infused 15-30 minutes after playback of the less-preferred song.

### Anatomy

Following the completion of experiments, birds were deeply anesthetized with isoflurane before being transcardially perfused with 0.9% saline, followed by 4% paraformaldehyde in 0.025 M phosphate buffer (PB). Brains were post-fixed for 4-hrs, then cryoprotected in 30% sucrose. Forty-micron parasagittal sections were cut on a freezing microtome and every third section was stained with cresyl violet acetate to determine the locations of cannulae and probes. All females in this study were confirmed to have probes located within NCM.

### Song stimuli

All male song stimuli were female-directed courtship song samples recorded from males (N=10) from our colony at McGill University. Songs were recorded in sound-attenuating chambers (TRA Acoustics, Cornwall, Ontario) by briefly exposing males to stimulus females (not used in this experiment), as has been previously described^22,26,58^. We created stimuli for five pairs of males for use in two-choice female preference tests. Pairs consisted of the same two males for all females (e.g. Male A and Male B were always pair 1, Male C and Male D were always pair 2, etc.), and females were tested on 2-5 pairs. Males used to generate song stimuli and females used in this study were unrelated and had never physically interacted. Song stimuli used for the preference test were matched for duration and number of introductory notes and were free of noise and female calls. For each male, we used one song example containing multiple motifs and introductory notes. All stimulus songs were bandpass filtered (300–10 kHz), normalized by their maximum amplitude, and saved as wav files (44.1 kHz) using custom written code in Matlab (Mathworks, Natick, MA). We used a selection of 4-7 recordings of song to provide a representative sample of varying song duration, number of bouts, and number of introductory notes from each male’s repertoire.

### Analyses

All statistical analyses were completed using JMP Statistical Processing Software (SAS, Cary, NC, USA) or custom-written Matlab code (Mathworks, Natick, MA). To quantify female preference for one song over the other, we determined the distribution of string pulls for Male A versus Male B. The initially preferred male was attributed a value of 1, and the less-preferred male a value of zero, and the distribution of pulls was bootstrapped with replacement (10,000 iterations) to obtain 95% confidence intervals. We also calculated the “strength of preference” (the distance of the bootstrapped distribution mean from 0.5, a chance distribution). This allowed us to separate the strength of preference from the directionality, for example if females had strong preferences overall but differed in which male they preferred.

To assess whether females demonstrated song preferences during the preference tests, we conducted two-tailed single-sample t-tests on mean “strength of preference” for each pair of males and tested whether the distribution of preference significantly differed from chance (H0=0.5). We also tested whether mean preference between the two males was skewed in any of our pairs by attributing one male a value of one and the other male a value of zero across all females. We then conducted two-tailed single-sample t-tests by pair to see whether the mean preference between two males was significantly different from 0.5 (chance).

To examine whether the drug manipulation paired with playback of the less-preferred male’s song was related to change in preference between Day 1 and Day 4, we performed paired t-tests of the bootstrapped means for Day 1 and Day 4 for each drug. We also tested whether the amount by which behavior shifted was related to drug manipulation using a model with percent change in string pull distribution as a dependent variable, drug as an independent variable, and female ID as a random variable. All models were conducted using a restricted maximum likelihood approach with unbounded variance components.

Finally, we examined whether overall changes in activity could account for the changes in preference. To do this, we used a model with percent change in string pull distribution as the dependent variable, percent change in overall string pulling as an independent variable, and bird ID as a random variable, independently for each drug.

## ACKNOWLEDGEMENTS

We would like to thank Jon T. Sakata for helpful discussions and comments on the manuscript. In addition, we would like to thank Therese Koch for help with preference testing and histology. This work was funded by the Natural Sciences and Engineering Research Council of Canada (NSERC) and Fonds de Recherche du Québec – Nature et technologies (FQRNT) to SCW, a McGill Integrated Program in Neuroscience returning student award to HJB, and a Lloyd Carr-Harris fellowship to EMW.

## AUTHOR CONTRIBUTIONS

HJB and SCW designed the experiments; HJB and EMW performed the experiments; HJB and SCW analyzed the data; HJB, EMW, and SCW wrote and edited the manuscript.

---

**Suppl. Figure 1.**
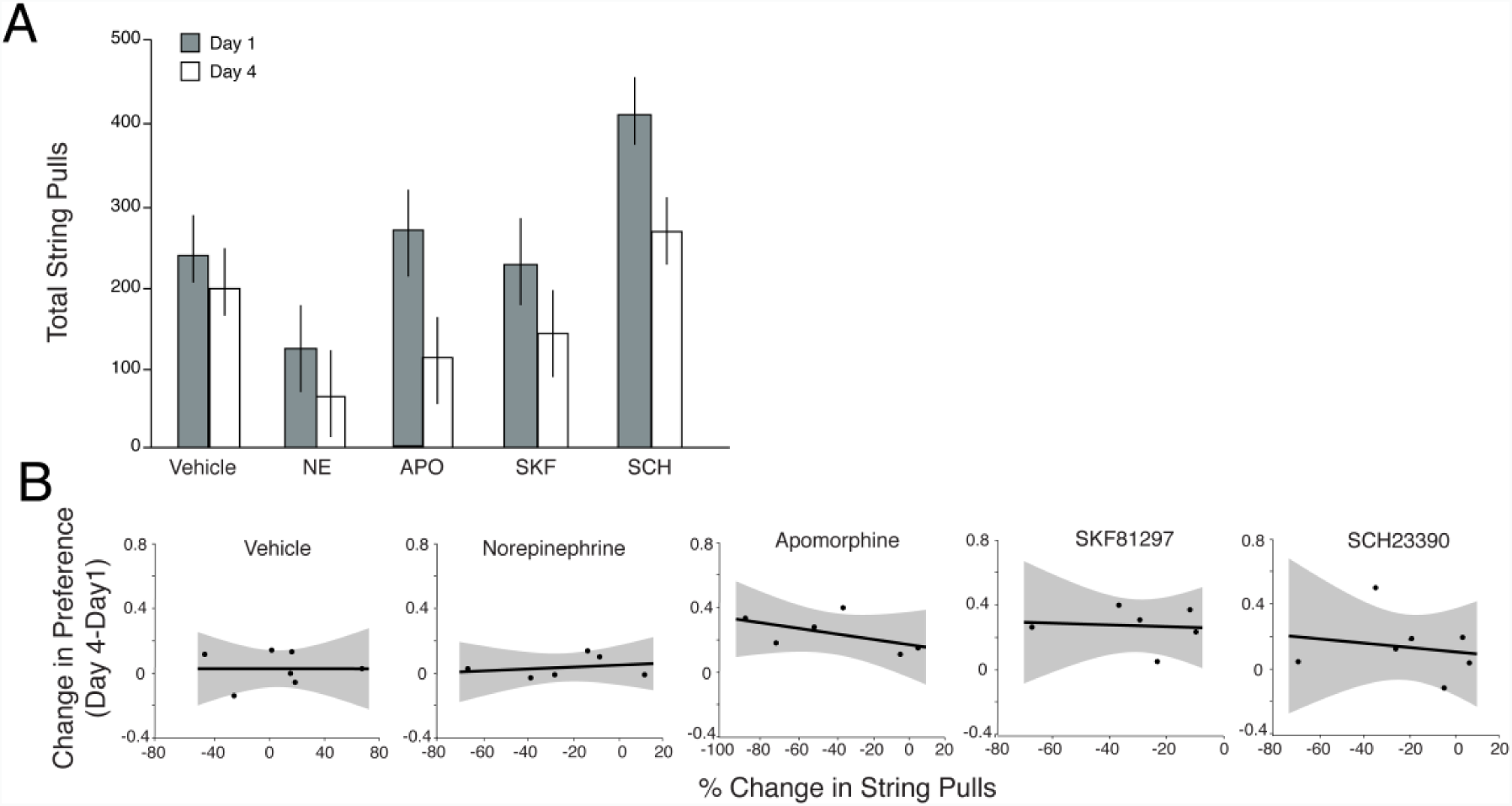
Preference changes not correlated with general changes in motor behavior or motivation. A) Across all conditions, the total number of string pulls decreased between Day 1 and Day 4. Vehicle (VEH), norepinephrine (NE), apomorphine (APO), D1 agonist SKF81297 (SKF), D1 antagonist SCH23390 (SCH). B) However, the percent change in string pulls was not significantly correlated with the change in preference for any of the conditions.

## REFERENCES

1. Bakin, J. S., South, D. A. & Weinberger N. M. Induction of receptive field plasticity in the auditory cortex of the guinea pig during instrumental avoidance conditioning. Behavioral Neuroscience 110, 905–913 (1996).

2. Edeline, J.-M., Pham, P. & Weinberger N. M. Rapid development of learning-induced receptive field plasticity in the auditory cortex. Behavioral Neuroscience 107, 539–551 (1993).

3. Froemke, R. C. & Jones B. J. Development of auditory cortical synaptic receptive fields. Neuroscience & Biobehavioral Reviews 35, 2105–2113 (2011).

4. Pienkowski, M. & Eggermont J. J. Cortical tonotopic map plasticity and behavior. Neuroscience & Biobehavioral Reviews 35, 2117–2128 (2011).

5. Miranda, J. A. & Wilczynski W. Female reproductive state influences the auditory midbrain response. J Comp Physiol A 195, 341–349 (2009).

6. Sisneros, J. A. Steroid-dependent auditory plasticity for the enhancement of acoustic communication: Recent insights from a vocal teleost fish. Hearing Research 252, 9–14 (2009).

7. Caras, M. L., Sen, K., Rubel, E. W. & Brenowitz E. A. Seasonal Plasticity of Precise Spike Timing in the Avian Auditory System. J. Neurosci. 35, 3431–3445 (2015).

8. Miranda, J. A. & Liu R. C. Dissecting natural sensory plasticity: Hormones and experience in a maternal context. Hearing Research 252, 21–28 (2009).

9. Marlin, B. J., Mitre, M., D’amour, J. A., Chao, M. V. & Froemke R. C. Oxytocin enables maternal behaviour by balancing cortical inhibition. Nature 520, 499–504 (2015).

10. Marlin, B. J. & Froemke R. C. Oxytocin modulation of neural circuits for social behavior. Developmental Neurobiology 77, 169–189 (2017).

11. Maney, D. L. The incentive salience of courtship vocalizations: Hormone-mediated ‘wanting’ in the auditory system. Hearing Research 305, 19–30 (2013).

12. Berridge, K. C. The debate over dopamine’s role in reward: the case for incentive salience. Psychopharmacology 191, 391–431 (2007).

13. Berke, J. D. What does dopamine mean? Nature Neuroscience 21, 787–793 (2018).

14. Schultz, W., Dayan, P. & Montague, P. R. A Neural Substrate of Prediction and Reward. Science 275, 1593–1599 (1997).

15. Bromberg-Martin, E. S., Matsumoto, M. & Hikosaka O. Dopamine in Motivational Control: Rewarding, Aversive, and Alerting. Neuron 68, 815–834 (2010).

16. Fields, H. L., Hjelmstad, G. O., Margolis, E. B. & Nicola S. M. Ventral Tegmental Area Neurons in Learned Appetitive Behavior and Positive Reinforcement. Annual Review of Neuroscience 30, 289–316 (2007).

17. Aragona, B. J., Liu, Y., Curtis, J. T., Stephan, F. K. & Wang, Z. A Critical Role for Nucleus Accumbens Dopamine in Partner-Preference Formation in Male Prairie Voles. J. Neurosci. 23, 3483–3490 (2003).

18. Gobrogge, K. & Wang Z. The ties that bond: neurochemistry of attachment in voles. Current Opinion in Neurobiology 38, 80–88 (2016).

19. Bao, S., Chan, V. T. & Merzenich M. M. Cortical remodelling induced by activity of ventral tegmental dopamine neurons. Nature 412, 79 (2001).

20. Puschmann, S., Brechmann, A. & Thiel C. M. Learning-dependent plasticity in human auditory cortex during appetitive operant conditioning. Human Brain Mapping 34, 2841–2851 (2013).

21. Salimpoor, V. N., Zald, D. H., Zatorre, R. J., Dagher, A. & McIntosh, A. R. Predictions and the brain: how musical sounds become rewarding. Trends in Cognitive Sciences 19, 86–91 (2015).

22. Schubloom, H. E. & Woolley S. C. Variation in social relationships relates to song preferences and EGR1 expression in a female songbird. Developmental Neurobiology 76, 1029–1040 (2016).

23. Riebel, K., Smallegange, I. M., Terpstra, N. J. & Bolhuis J. J. Sexual equality in zebra finch song preference: evidence for a dissociation between song recognition and production learning. Proceedings of the Royal Society of London. Series B: Biological Sciences 269, 729–733 (2002).

24. Barr, H. J. & Woolley S. C. Developmental auditory exposure shapes responses of catecholaminergic neurons to socially-modulated song. Scientific Reports 8, (2018).

25. Bolhuis, J. J. & Gahr M. Neural mechanisms of birdsong memory. Nature Reviews Neuroscience 7, 347–357 (2006).

26. Woolley, S. C. & Doupe A. J. Social Context–Induced Song Variation Affects Female Behavior and Gene Expression. PLOS Biology 6, e62 (2008).

27. Yanagihara, S. & Yazaki-Sugiyama, Y. Auditory experience-dependent cortical circuit shaping for memory formation in bird song learning. Nature Communications 7, 11946 (2016).

28. Chew, S. J., Vicario, D. S. & Nottebohm, F. A large-capacity memory system that recognizes the calls and songs of individual birds. Proceedings of the National Academy of Sciences 93, 1950–1955 (1996).

29. Kubikova, L., Wada, K. & Jarvis E. D. Dopamine receptors in a songbird brain. The Journal of Comparative Neurology 518, 741–769 (2010).

30. Lou, Y. et al. Ventral tegmental area activation promotes firing precision and strength through circuit inhibition in the primary auditory cortex. Front. Neural Circuits 8, (2014).

31. Shepard, K. N., Kilgard, M. P. & Liu R. C. Experience-Dependent Plasticity and Auditory Cortex. in Neural Correlates of Auditory Cognition (eds. Cohen, Y. E., Popper, A. N. & Fay, R. R.) 293–327 (Springer New York, 2013). doi:10.1007/978-1-4614-2350-8_10

32. Reed, A. et al. Cortical Map Plasticity Improves Learning but Is Not Necessary for Improved Performance. Neuron 70, 121–131 (2011).

33. Carcea, I. & Froemke R. C. Chapter 3 - Cortical Plasticity, Excitatory–Inhibitory Balance, and Sensory Perception. in Progress in Brain Research (eds. Merzenich, M. M., Nahum, M. & Van Vleet, T. M.) 207, 65–90 (Elsevier, 2013).

34. Matragrano, L. L. et al. Rapid Effects of Hearing Song on Catecholaminergic Activity in the Songbird Auditory Pathway. PLoS ONE 7, e39388 (2012).

35. Tanaka, M., Sun, F., Li, Y. & Mooney, R. A mesocortical dopamine circuit enables the cultural transmission of vocal behaviour. Nature 563, 117–120 (2018).

36. Hamid, A. A. et al. Mesolimbic dopamine signals the value of work. Nature Neuroscience 19, 117–126 (2016).

37. Howe, M. W. & Dombeck D. A. Rapid signalling in distinct dopaminergic axons during locomotion and reward. Nature 535, 505–510 (2016).

38. Ding, L. & Perkel D. J. Long-Term Potentiation in an Avian Basal Ganglia Nucleus Essential for Vocal Learning. J. Neurosci. 24, 488–494 (2004).

39. Ding, L. & Perkel D. J. Dopamine Modulates Excitability of Spiny Neurons in the Avian Basal Ganglia. J. Neurosci. 22, 5210–5218 (2002).

40. Calabresi, P., Picconi, B., Tozzi, A. & Di Filippo, M. Dopamine-mediated regulation of corticostriatal synaptic plasticity. Trends in Neurosciences 30, 211–219 (2007).

41. Otani, S., Daniel, H., Roisin, M.-P. & Crepel F. Dopaminergic Modulation of Long-term Synaptic Plasticity in Rat Prefrontal Neurons. Cereb Cortex 13, 1251–1256 (2003).

42. Sheynikhovich, D., Otani, S. & Arleo A. Dopaminergic Control of Long-Term Depression/Long-Term Potentiation Threshold in Prefrontal Cortex. J. Neurosci. 33, 13914–13926 (2013).

43. Wise, R. A. Dopamine, learning and motivation. Nature Reviews Neuroscience 5, 483–494 (2004).

44. Lisman, J., Grace, A. A. & Duzel, E. A neoHebbian framework for episodic memory; role of dopamine-dependent late LTP. Trends in Neurosciences 34, 536–547 (2011).

45. London, S. E. & Clayton D. F. Functional identification of sensory mechanisms required for developmental song learning. Nature Neuroscience 11, 579–586 (2008).

46. Gobes, S. M. H. & Bolhuis J. J. Birdsong Memory: A Neural Dissociation between Song Recognition and Production. Current Biology 17, 789–793 (2007).

47. Miller, D. B. The acoustic basis of mate recognition by female zebra finches. anim behav 27, 376–380 (1979).

48. Miller, D. B. Long-term recognition of father’s song by female zebra finches. Nature 280, 389–391 (1979).

49. Clayton, N. S. Song discrimination learning in zebra finches. Animal Behaviour 36, 1016–1024 (1988).

50. Happel, M. F. K. Dopaminergic impact on local and global cortical circuit processing during learning. Behavioural Brain Research 299, 32–41 (2016).

51. Sara, S. J. & Bouret S. Orienting and Reorienting: The Locus Coeruleus Mediates Cognition through Arousal. Neuron 76, 130–141 (2012).

52. Acerbo, M. J. & Delius J. D. Behavioral Sensitization to Apomorphine in Pigeons (Columba livia): Blockade by the D_1_ Dopamine Antagonist SCH-23390. Behavioral Neuroscience 118, 1080–1088 (2004).

53. Rose, J., Schiffer, A.-M., Dittrich, L. & Güntürkün, O. The role of dopamine in maintenance and distractability of attention in the “prefrontal cortex” of pigeons. Neuroscience 167, 232–237 (2010).

54. Herold, C., Diekamp, B. & Güntürkün, O. Stimulation of dopamine D1 receptors in the avian fronto-striatal system adjusts daily cognitive fluctuations. Behavioural Brain Research 194, 223–229 (2008).

55. Diekamp, B., Kalt, T., Ruhm, A., Koch, M. & Güntürkün, O. Impairment in a discrimination reversal task after D1 receptor blockade in the pigeon ‘prefrontal cortex’. Behavioral Neuroscience 114, 1145–1155 (2000).

56. Ikeda, M. Z., Jeon, S. D., Cowell, R. A. & Remage-Healey, L. Norepinephrine Modulates Coding of Complex Vocalizations in the Songbird Auditory Cortex Independent of Local Neuroestrogen Synthesis. Journal of Neuroscience 35, 9356–9368 (2015).

57. Tchernichovski, O., Nottebohm, F., Ho, C. E., Pesaran, B. & Mitra, P. P. A procedure for an automated measurement of song similarity. Animal Behaviour 59, 1167–1176 (2000).

58. Chen, Y., Clark, O. & Woolley S. C. Courtship song preferences in female zebra finches are shaped by developmental auditory experience. Proceedings of the Royal Society B: Biological Sciences 284, 20170054 (2017).

